# RNase H1 levels dramatically affect mitochondrial genome maintenance with little impact on nuclear R-loops in murine B cells

**DOI:** 10.1101/2025.04.30.651504

**Authors:** Kiran Sakhuja, Stella R Hartono, Lionel A Sanz, Eriko Ijiri, Caitlin Darling, Louis Dye, James R. Iben, Hyongi Chon, Susana Cerritelli, Robert Crouch, Frédéric Chédin

**Affiliations:** Division of Intramural Research, Eunice Kennedy Shriver National Institute of Child Health and Human Development, National Institutes of Health, Bethesda, MD, USA; Department of Molecular and Cellular Biology, University of California, Davis, CA 95616, USA; Microscopy and Imaging Core, Eunice Kennedy Shriver National Institute of Child Health and Human Development, National Institutes of Health, Bethesda, MD, USA; Mouse Genetics Core, Eunice Kennedy Shriver National Institute of Child Health and Human Development, National Institutes of Health, Bethesda, MD, USA

## Abstract

Overexpression of RNase H1, a ribonuclease that degrades RNA:DNA hybrids and R-loops, can suppress genome instability phenotypes in a range of maladaptive conditions. This has been interpreted to suggest that genotoxic co-transcriptional R-loops arise under these conditions and are resolved by RNase H1. Here, we manipulated RNase H1 levels using conditional knockout and overexpression models in primary murine B cells and mapped the resulting genomic R-loop landscapes. *Rnaseh1* deletion resulted in a dramatic loss of mitochondrial replication and compromised B cell responses, consistent with a critical mitochondrial function for RNase H1. Genome-wide R-loops were, however, not significantly affected. More surprisingly, overexpressing active nuclear RNase H1 did not lead to significant reduction of R-loop levels or change their distribution. These results were confirmed using a human cell line in which active, nuclear RNase H1 can be induced. Our findings indicate that co-transcriptional R-loops are not efficiently resolved by RNase H1 and suggest that the identity of the RNA/DNA hybrids at the root of the genome instability phenotypes suppressed by RNase H1 may need to be re-interpreted.

## INTRODUCTION

Ribonucleases H (RNases H) are highly conserved enzymes that cleave the RNA strand of RNA/DNA hybrids (Stein & Hausen, 1969). Two RNases H (H1 and H2) are present in most organisms and have the possibility of sharing substrates, although they also have non-overlapping substrates (Cerritelli & Crouch, 2009). We previously observed that RNase H1 localizes to both nuclei and mitochondria, where it plays a critical role in mitochondrial DNA (mtDNA) replication (Cerritelli *et al*, 2003; Holmes *et al*, 2015). RNase H2, by contrast, is exclusively nuclear, and possesses a unique role in removing ribonucleotides (rNMPs) incorporated by DNA polymerases during replication of genomic DNA (McElhinny *et al*, 2010; Sparks *et al*, 2012). Both RNases H are essential for embryonic development. In mouse, deletion of the *Rnaseh1* gene or deletion of any of the genes encoding any of the subunits of the heterotrimeric RNase H2 protein, results in embryonic arrest around embryonic day 9 (Cerritelli *et al*., 2003; Reijns *et al*, 2012). Hypomorphic mutations in human RNase H1 result in muscle weakness (Akman *et al*, 2016; Bugiardini *et al*, 2017; Reyes *et al*, 2015), typically observed in mitochondrial dysfunctions, while hypomorphic mutations in any of the three genes encoding the heterotrimeric RNase H2 protein induce Aicardi-Goutières syndrome, a type I interferonopathy (Crow *et al*, 2006; Crow & Manel, 2015).

RNA/DNA hybrids are biochemical substrates for both types of RNase H, and are found in cells in the form of R-loops, a type of three-stranded nucleic acid structure composed of an RNA/DNA hybrid and a displaced DNA strand (Thomas *et al*, 1976), are biochemical substrates for both types of RNase H. R-loops are thought to form during transcription upon the reannealing of the nascent RNA with the template DNA strand. In mammalian cells, R-loops were first shown to form over the class switch recombination (CSR) regions involved in immunoglobulin isotype switching in B cells (Daniels & Lieber, 1995; Reaban & Griffin, 1990; Reaban *et al*, 1994; Yu *et al*, 2003). The role of R-loops in CSR remains debated; a B cell-specific RNase H1 overexpression mouse model did not lead to reduced CSR (Maul *et al*, 2017). The locations of R-loops in the genome can be inferred using the S9.6 anti-RNA/DNA hybrid antibody by a methodology termed DNA/RNA Immunoprecipitation (DRIP-seq). DRIP-based methods have revealed that R-loops are prevalent, dynamic structures that can form at appreciable frequencies over thousands of transcribed genomic loci in mammalian cells under normal conditions (Chedin, 2016; Crossley *et al*, 2020; Ginno *et al*, 2012; Sanz *et al*, 2016). Work in Bacteria, yeasts, worms, flies, and plants has further suggested that R-loop formation is universal during transcription.

Deregulated R-loop homeostasis has often been invoked as a cause for genome instability phenotypes observed under a variety of pathological conditions, particularly those associated with dysfunctional RNA metabolism (Crossley *et al*, 2019; Garcia-Muse & Aguilera, 2019). The suppression of genome instability phenotypes via RNase H1 overexpression in cells has become the gold standard for defining phenomena associated with “harmful” R-loops (Cerritelli *et al*, 2022; Huertas & Aguilera, 2003). Few studies, however, have directly shown that RNase H1 overexpression indeed led to a significant reduction of harmful genomic R-loops.

Expression of a catalytically inactive RNase H1 protein in cells has also been used to determine the chromosomal location of R-loops (RChIP-seq) (Chen *et al*, 2017). This methodology confirms prevalent R-loop formation, although RNase H1-based and S9.6 antibody-based R-loop maps show only partial overlaps (Castillo-Guzman & Chedin, 2021; Miller *et al*, 2022). RNase H1-based maps are biased towards promoter and enhancer regions associated with paused RNA polymerase (Chen *et al*., 2017), while S9.6-based maps provide broader coverage, identifying transcribed regions associated with elongating RNA polymerase as primary targets. This raises the possibility that distinct sub-classes of R-loops or RNA/DNA hybrids exist and that RNase H1 may have differential access to these sub-classes due to either selective targeting or restricted access mechanisms. Thus, our understanding of the *in vivo* genomic substrates of RNase H1 is incomplete.

Here, we manipulated RNase H1 expression levels using both a conditional *Rnaseh1* knockout, and a conditional RNase H1 overexpression model specifically in resting and stimulated splenic murine B cells. R-loop distribution was then assessed genome-wide in these primary cells using S9.6 antibody-based approaches. We confirm that loss of *Rnaseh1* in B cells leads to dramatic mitochondrial phenotypes and compromised B cell responses to *ex vivo* stimulation. Genome-wide R-loops, however, were not significantly affected. More surprisingly, RNase H1 overexpression in B cells did not lead to significant reduction of nuclear R-loop loads.

## RESULTS

### A mouse model for conditional *Rnaseh1* deletion in B cells

We previously described murine embryonic fibroblasts (MEFs) from a mouse strain in which the *Rnaseh1* gene can be conditionally knocked out in response to Cre recombinase expression (Holmes *et al*., 2015). In the present study, we chose to use mouse B cell development as a rigorous and well-characterized model to examine the consequences of loss of RNase H1 expression in animals. Cre expression was driven by the *mb1* promoter to conditionally delete a portion of the *Rnaseh1* gene including exons 5 to 7 (Figure 1A). The *mb1* gene encodes the Ig-α signaling subunit of the B cell antigen receptor, which is activated at the Early Pro-B stage and continuously expressed in the B cell lineage in all later stages except in plasma cells (Hobeika *et al*, 2006). Mice with loxP sites flanking exons 5-7 of the *Rnaseh1* gene were crossed with a mouse expressing EIIA-Cre, obtaining germline deletion transmission. *Rnaseh1* ^+/del^ mice were then crossed with mice bearing the floxed allele of *Rnaseh1^fl/fl^* and screened to obtain *Rnaseh1* ^fl/del^ or conditional / deletion mice. Hereafter, we will use *H1* ^fl/del^ for the control strain and *H1* ^del/del^ for the strain expressing the *mb1*-driven Cre in which the floxed *RNaseh1* allele is deleted in early-Pro-B cells, leading to a conditional knockout.

**Figure 1:**
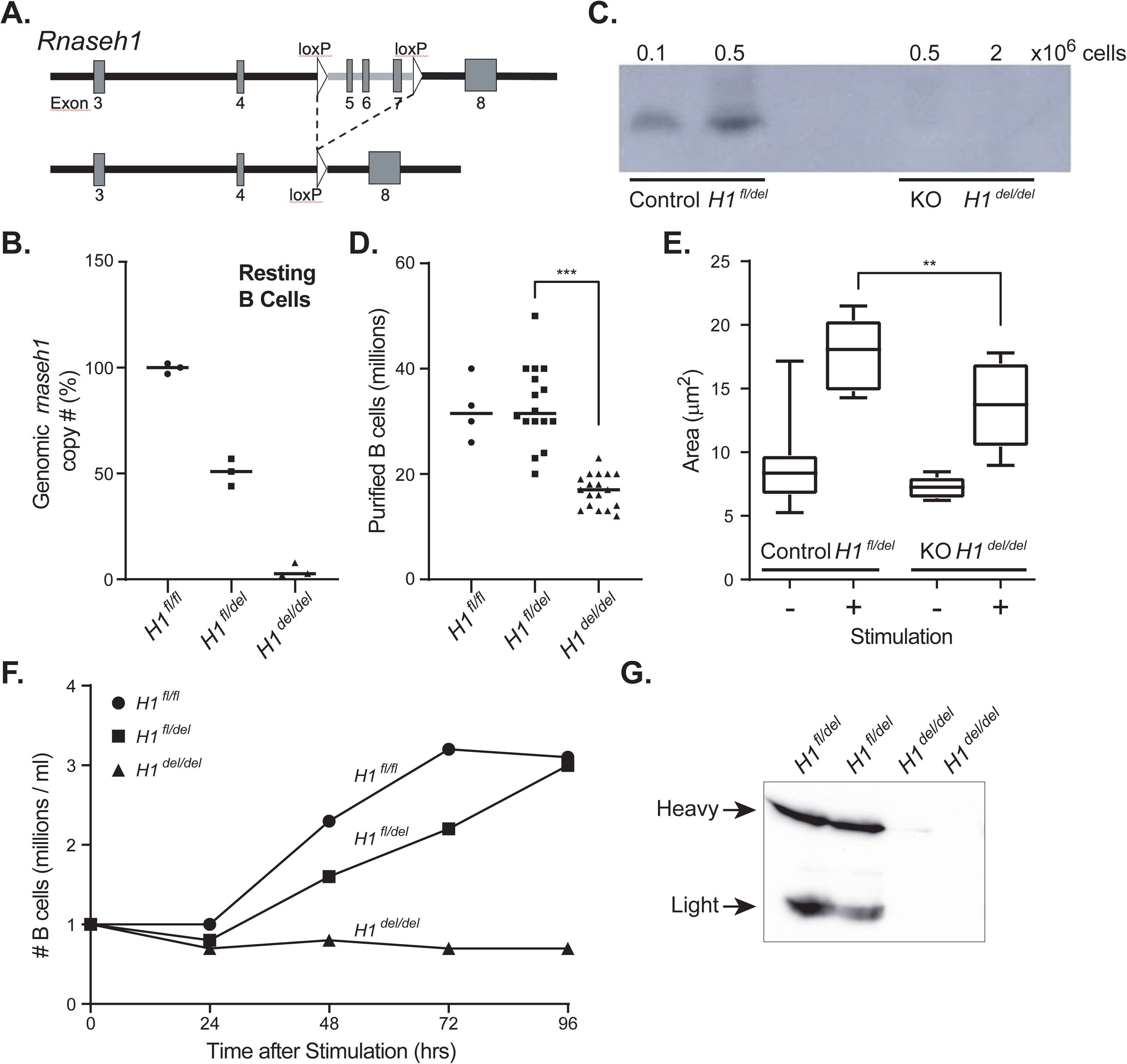
**A.** Schematic description of a portion of the murine *Rnaseh1* gene with the position of the loxP sites indicated. Cre expression leads to the conditional deletion of exons 5-7. **B**. Relative copy number of the floxed region in the genome of resting B cells of the indicated genotypes. **C**. Gel-based RNase H1 activity assay. The ability of endogenous RNase H1 to cleave labeled RNA:DNA hybrid substrates is indicated by the presence of a band. The number of B cells used to produce protein lysates carrying RNase H1 is indicated at the top. Corresponding genotypes are at bottom. **D**. Number of resting B cells purified per mouse spleen of the corresponding genotypes; each symbol corresponds to one animal. **E**. Cell area of resting and stimulated B cells measured by transmission electron microscopy, displayed as boxplots for each corresponding genotype. **F**. B cell proliferation following stimulation of 1 million cells is depicted as the number of cells over time post stimulation for each genotype. **G**. Representative western blot depicting the production of antibodies in sera from animals of the corresponding genotypes.

### *Rnaseh1* deletion leads to altered B cell development and proliferation

We characterized resting splenic B cells in *H1* ^del/del^ and *H1* ^fl/del^ strains and verified that H1 ^del/del^ B cells had almost no nuclear *Rnaseh1* DNA within the deleted region, while H1 ^fl/del^ cells had one allele and control animals, two alleles (Figure 1B). As expected, H1 ^del/del^ B cells displayed no detectable RNase H1 enzymatic activity against labeled RNA/DNA hybrid substrates, confirming functional gene knockout (Figure 1C). H1 ^del/del^ animals produced 50% fewer resting B cells than the corresponding H1 ^fl/del^ strain (Figure 1D; p<0.0001, unpaired t-test), suggesting a proliferation or developmental issue. We next examined the response of naïve B cells to stimulation with lipopolysaccharide (LPS) and interleukin-4 (Il-4) for 24 h. As expected, resting B cells from H1 ^fl/del^ and H1 ^del/del^ animals were small with little cytoplasmic volume. Stimulation resulted in significant increases in nuclear volume for both genotypes, consistent with B cell swelling upon stimulation. However, the cytoplasmic area of H1 ^fl/del^ cells was significantly greater (p=0.0136, unpaired t-test) than the corresponding area from *Rnaseh1*-deficient B cells (Figure 1E), suggesting a compromised ability to respond to stimulation. Further analysis showed that H1 ^del/del^ B cells were defective in proliferation assays (Figure 1F) and cell cycle analysis revealed that they were arrested at the G1/S phase. Finally, sera collected from H1 ^del/del^ mice had very little, if any, circulating antibodies (Figure 1G), indicating a failure to initiate the adaptive immune response and produce antibodies. Our results suggest that loss of RNase H1 activity dramatically compromises B cell development and the induction of the B cell program upon ex vivo stimulation.

### RNase H1 loss impacts gene expression patterns, supporting a decreased proliferative potential

We performed RNA-seq analysis of resting and stimulated B cells from both strains to capture the impact of RNase H1 loss on gene expression programs. When compared to resting control cells, stimulated control cells showed vast transcriptome changes, consistent with prior observations (Figure S1A, Table S1) (Kieffer-Kwon *et al*, 2017; Kouzine *et al*, 2013). Many up-regulated genes were involved in functions related to cell cycle, DNA repair, RNA metabolism, ribosome biogenesis and mitochondrial biology, consistent with the rapid induction of many pro-growth genes and the resumption of cell proliferation and active metabolism (Figure S1B). Similar gene ontology (GO) enrichments were observed when analyzing nascent transcription patterns using EU-seq, with a significant overlap between RNA-seq and EU-seq datasets, as expected (Figure S1C). Expression gene analysis also showed that H1 ^del/del^ cells underwent strong induction of the B cell differentiation program upon stimulation, with similar gene classes showing up- and down-regulation (Figure 2A). Thus, RNase H1 depletion did not abrogate the gene expression response of naïve B cells to stimulation.

**Figure 2:**
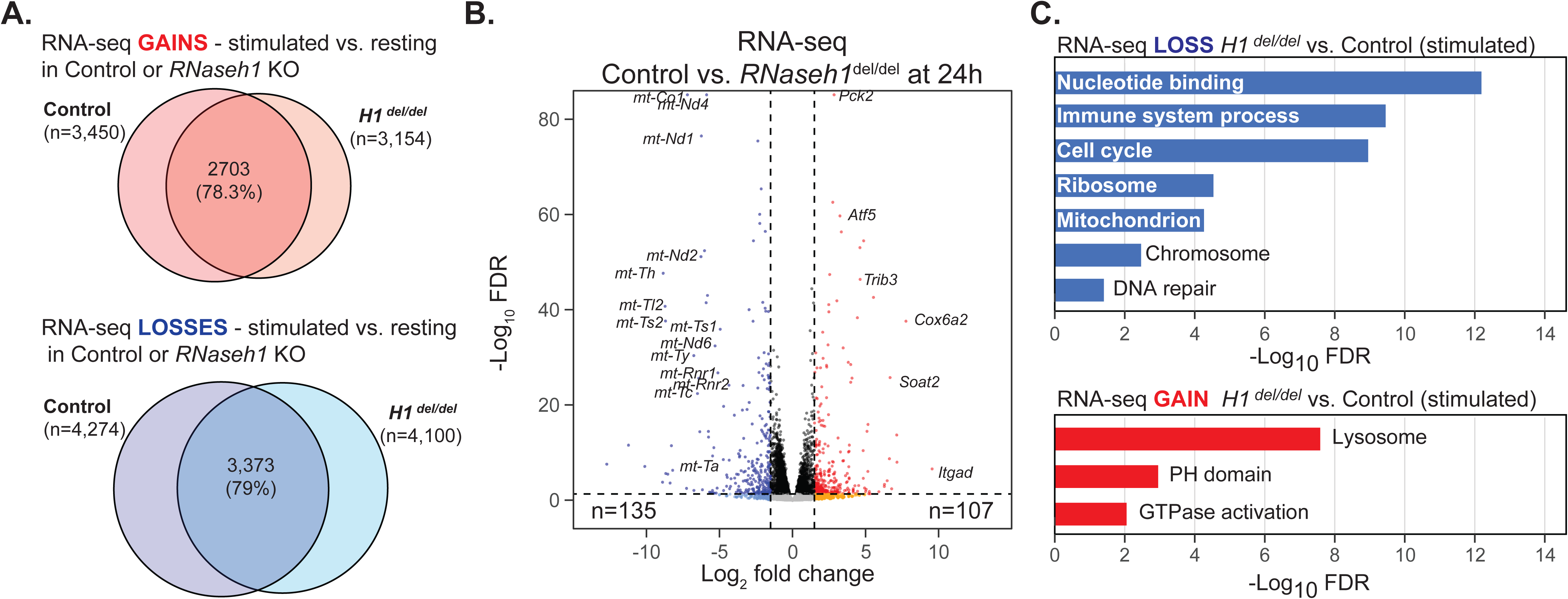
**A.** Venn diagrams depicting overlap between genes that were upregulated (Gain) or downregulated (Loss) in control cells (*H1* ^fl/del^) and *Rnaseh1* KO cells (*H1* ^del/del^) upon B cell stimulation. **B.** Volcano plot of RNA-seq data comparing gene expression differences between *Rnaseh1* KO cells (*H1* ^del/del^) and control cells (*H1* ^fl/del^) post stimulation. Vertical dashed lines indicate |Log_2_FC|>1 and the horizontal dashed line indicates a Log_10_(FDR)<0.05. Relevant genes are indicated along with the total number of significantly up- or down-regulated genes. **C.** Significantly enriched Gene Ontologies (David) are indicated for down- and up-regulated genes comparing *Rnaseh1* KO cells (*H1* ^del/del^) and control cells (*H1* ^fl/del^) post stimulation.

Comparison of RNA-seq datasets between H1 ^del/del^ and H1 ^fl/del^ cells 24 h post stimulation nonetheless revealed significant differences in gene expression states (Figure 2B). Nearly 2,000 genes were significantly down-regulated in *Rnaseh1* KO cells with clear GO enrichment for genes involved in pro-growth pathways such as genes involved in cell cycle, DNA replication, mitochondria, ribosome biogenesis and immune system (Figure 2C). About 1,200 genes also showed significant up-regulation including genes involved in lysosomal biology and signal transduction (Figure 2C). Thus, while H1 ^del/del^ B cells broadly attempt to initiate a regular transcriptional response to B cell stimulation, it appears quantitatively different and biased towards a lower proliferative potential.

To better understand the impact of RNase H1 loss on the transcriptional state of H1 ^del/del^ B cells post stimulation, we focused on 125 genes listed as cell cycle genes in the Kegg database. 81 (65%) genes showed no significant changes in expression between H1 ^del/del^ and H1 ^fl/del^ genotypes at the resting stage, confirming the general G0 state of resting H1 ^del/del^ cells. However, expression of twelve cell cycle genes was significantly changed between KO and control cells at the resting stage: ten increased, two decreased. *Ccnd2* and *Cdk6*, which are both important for the G1/S cell cycle transition (Martinez-Alonso & Malumbres, 2020) were the most up-regulated genes at the resting stage (Figure S1D). The two downregulated genes were *Gadd45b*, which is related to DNA damage during G2 to M transition (Magimaidas *et al*, 2016), and *Cdkn1b* (p27Kip1), which is known to inhibit entry into the cell cycle (Martinez-Alonso & Malumbres, 2020). This suggests that the *RNaseh1* KO cells are precociously primed for exit from G0. Post stimulation, 33 genes showed significant down-regulation, including multiple genes involved in DNA replication initiation such as six members of the *Mcm* family, *Orc1*, and *cdc6* genes (Table S1), and cell cycle control genes, including *Ccnd2* which was over-expressed at the resting stage. This supports the observation of decreased B cell proliferative potential upon stimulation when lacking RNase H1 activity.

### RNase H1 loss causes a profound loss of mitochondrial gene expression, mitochondrial DNA, and the induction of the mitochondrial unfolded protein response (UPR^mt^) pathway

Given prior evidence that RNase H1 is required for mitochondrial DNA replication in early embryonic development (Cerritelli *et al*., 2003), we focused our analysis on the expression of genes that code for proteins with strong evidence for localization in mitochondria, including genes carried by the mtDNA and nuclear genomes (Mitocarta genes). Mitochondrial genes were amongst the genes showing the most dramatic reduction in gene expression (Figure 2B). In resting B cells, mitochondrial genes showed a 5-18-fold lower expression in *H1* ^dl/dl^ cells compared to *H1* ^fl/del^ controls (Table S1, Figure 3A). This trend became even more pronounced (10-35-fold) post stimulation. Six protein-coding genes located contiguously on the mitochondrial genome between nucleotides 7,000 and approximately 10,250 had low or absent read counts. Overall, *Rnaseh1*-deleted cells showed about 10-fold less RNA-seq reads mapping to the mitochondrial genome compared to control cells (Figure 3B). This suggests that the loss of RNase H1 gravely compromises mitochondrial gene expression and function. We next asked if mtDNA loss could account for the dramatic reduction of mtDNA gene expression by quantifying mtDNA copy number using qPCR. Resting B cells from *H1* ^del/del^ mice carried less than five percent of normal mtDNA amounts (Figure 3C). While mtDNA copy number increased significantly in stimulated control B cells, *H1* ^del/del^ B cells mtDNA copy number remained at least one order of magnitude lower than control cells. Characterization of mitochondrial morphology using transmission electron microscopy revealed that, in addition, mitochondria from *H1* ^del/del^ B cells displayed abnormal morphology (Figure 3D). Mitochondria in both resting and stimulated *Rnaseh1*-deleted cells were hollowed out and showed much lower cristae formation, indicating disruption in organelle function. Thus, as previously described during embryonic development (Cerritelli *et al*., 2003) and in hepatocytes (Lima *et al*, 2016), RNase H1 activity is critical for mtDNA genome replication and maintenance in B cells. It is likely that the reduced proliferative potential and inability to produce antibodies observed in *Rnaseh1* KO cells harks back to a loss of mitochondrial potential.

**Figure 3:**
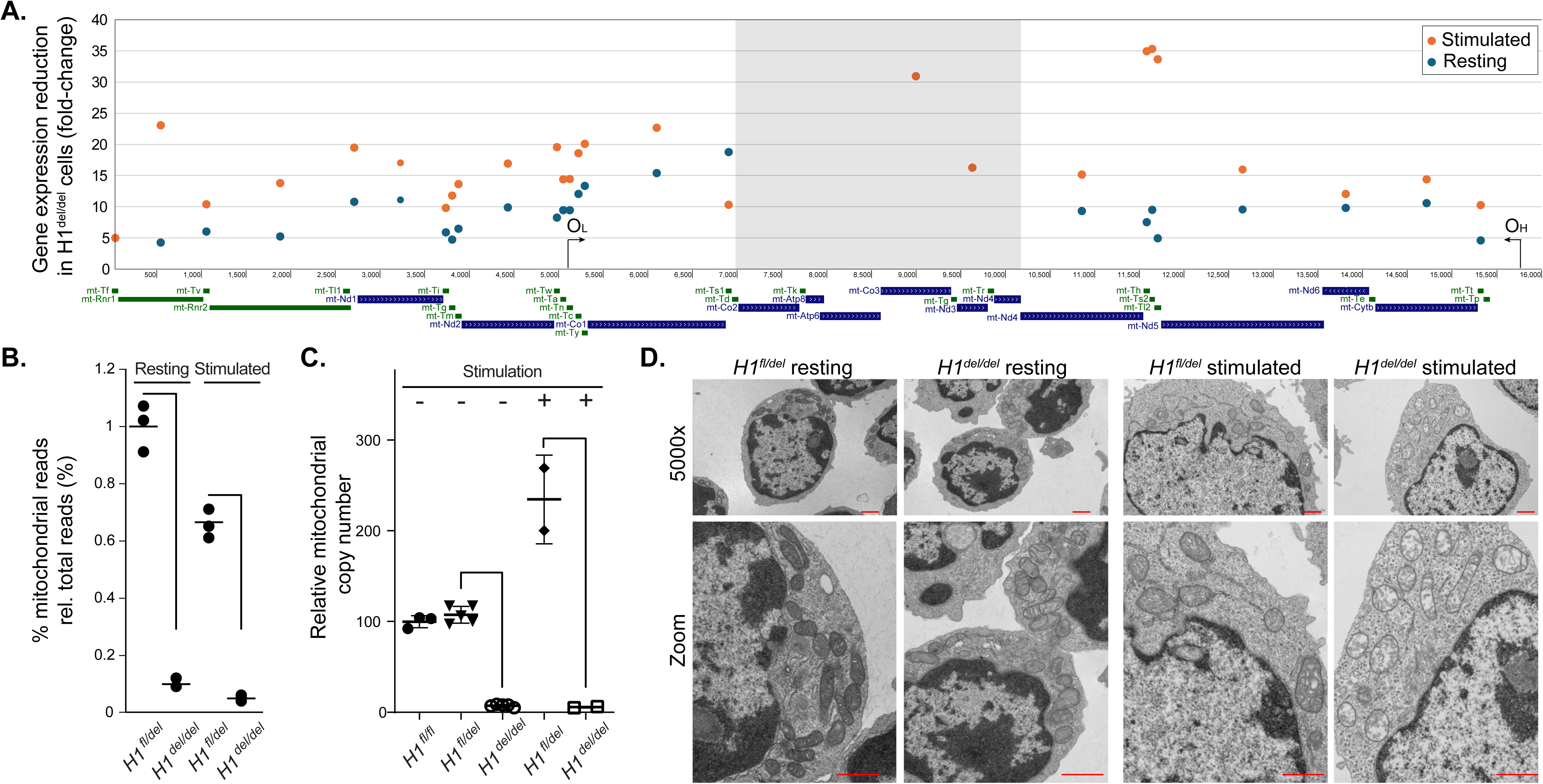
**A.** Gene expression defects are shown as an average fold-reduction between control cells and *Rnaseh1*-deleted cells for each gene along the mitochondrial genome. Fold-reduction was calculated from RNA-seq data and each data point is centered on its gene for both resting and stimulated conditions. The grey highlighted region has very few, if any read counts in KO cells. **B.** Quantification of total mitochondrial RNA-seq reads for *Rnaseh1*-deleted cells and control cells, pre- and post-stimulation. **C.** Analysis of relative mtDNA copy number for WT, control, and *Rnaseh1*-deleted cells pre- and post-stimulation. **D**. Transmission electron microscopy of resting and stimulated B cells from *Rnaseh1*-deleted and control samples. Scale bar is 1 micron. The zoomed area allows to focus on mitochondrial morphology.

RNA-seq data analysis also indicated that the loss of RNase H1 led to the up-regulation of the mitochondrial unfolded protein response (UPR^mt^) 24 hours post stimulation. The *Atf4* and *Ddit3* genes, encoding for key transcription factors that activate a common set of genes in response to various mitochondrial stresses (Shpilka & Haynes, 2018) were upregulated 1.7-fold and 2.3-fold, respectively in H1^del/del^ cells (Table S1). The *Atf5* gene encoding for a secondary UPR^mt^ transcription factor was up-regulated 5.2-fold, as were other downstream targets such as *Gdf15* (27-fold), *Soat2* (22-fold), *Cebpb* (3.2-fold), *Sesn2* (4.2-fold) and *Eif4ebp1* (2.5-fold) (Table S1). These measurements were validated by RT-qPCR for *Atf4*, *Atf5* and *Eif4ebp1* (Figure S2A) and confirmed by Western blots (Figure S2B). Overall, these results suggest that RNase H1 deficiency causes profound mitochondrial dysfunction and induction of the UPR^mt^.

### Loss of RNase H1 doesn’t significantly alter the nuclear R-loop landscape

To determine if loss of RNase H1 affected R-loop distribution, we mapped R-loops genome-wide using DRIP-seq and sDRIP-seq (Sanz *et al*, 2021) of *H1* ^fl/del^ and *H1* ^del/del^ B cells either at the resting stage or 24 hours post stimulation. As expected from the transcriptional effects observed in RNA-seq and EU-seq data, numerous loci showed significant R-loop gains and losses at thousands of loci upon stimulation in control cells (Figure S3A). When focusing on the heavy chain immunoglobulin region IgH, stimulation led to the expansion of the R-loop zone at the IgM / Sμ region, as best seen with the high-resolution sDRIP-seq method (Figure 4A). As expected from the LPS/IL-4 stimulation, sharp R-loop increases were also observed around the IgG1 / Sγ1, and to a lesser extent IgG3 / Sγ3, regions in control samples (Figure 4A). This is in agreement with prior locus-restricted approaches (Huang *et al*, 2007; Huang *et al*, 2006; Yu *et al*., 2003) and confirms that the IgH region is one of the genomic regions that undergoes high R-loop induction upon B cell stimulation. The R-loop increase was accompanied by a strong induction of nascent transcription measured by EU-seq and the appearance of the IgG1 mRNA transcript in RNA-seq data (Figure S3B). This suggests that R-loop mapping using DRIP-seq methods recapitulated expected key features of the B cell system, including a specific, powerful, R-loop increase around class-switch regions upon B cell stimulation.

**Figure 4:**
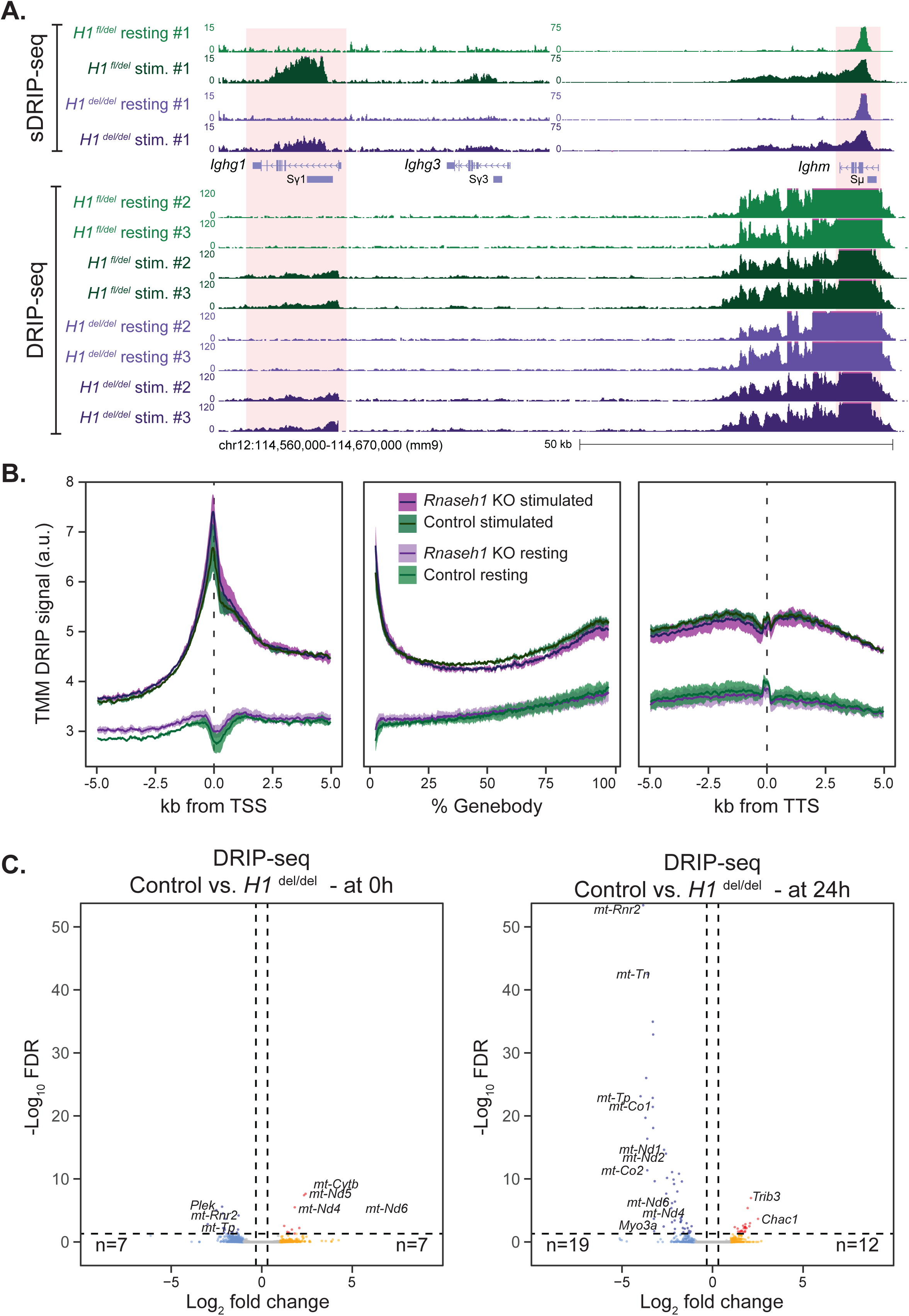
**A.** Genome browser screenshot showing R-loop distribution over the IgH region. High resolution sDRIP-seq datasets are shown at top for both control (*H1* ^fl/del^) and *Rnaseh1*-deficient (*H1* ^del/del^) samples; lower resolution DRIP-seq datasets for different animals are displayed at bottom. The IgM and IgG1 regions are highlighted. **B.** Metaplots of DRIP-seq signal are depicted over promoter, gene body, and terminal gene regions for control (*H1* ^fl/del^) and *Rnaseh1*-deficient (*H1* ^del/del^) samples with and without stimulation, as depicted. Line represents the trimmed mean while the 95% confidence interval is shown as a shaded band. **C**. Volcano plots of DRIP-seq data comparing R-loop intensities between control (*H1* ^fl/del^) and *Rnaseh1* KO cells (*H1* ^del/del^) at the resting (left) and stimulated (right) stages. Vertical dashed lines indicate |Log_2_FC|>1 and the horizontal dashed line indicates a Log_10_(FDR)<0.05.

R-loops were similarly induced at the IgM / Sμ and IgG1 / Sγ1 region in *Rnaseh1* KO cells (Figure 4A) and globally (Figure S3C, D), consistent with the notion that the transcriptional response to stimulation in *Rnaseh1*-deficient cells resembles that of control cells. However, R-loop increases were blunted compared to control conditions, consistent with evidence that naïve B cells lacking *Rnaseh1* are less able to mount a strong transcriptional response to stimulation (Figure 2). Remarkably, comparison of R-loop genic patterns between control and *Rnaseh1*-deficient samples revealed only few differences globally, whether in resting or stimulated B cells (Figure 4B, C). Post stimulation, the loci showing the largest R-loop losses corresponded to mitochondrially-encoded genes, as expected (Figure 4C). Overall, these data suggest that, at least in B cells, loss of RNase H1 does not significantly affect the R-loop landscape, indicating that RNase H1 does not play a major role in R-loop resolution in this cell type.

### Indirect transcriptomic changes in a B cell-specific transgenic nuclear RNase H1 overexpression model

To further assess the impact of nuclear RNase H1 on R-loop turnover, we took advantage of a transgenic mouse model expressing the nuclear form of *Rnaseh1* (devoid of the first 27 amino-acids) downstream of the H-2K^b^ promoter in a construct directly linked to the powerful B cell Eμ enhancer (Maul *et al*., 2017). In this setup, RNase H1 is overexpressed starting from pre-pro B cells and showed clear increase in protein abundance and a 50-100-fold increased activity against labeled RNA/DNA hybrids. In contrast to RNase H1 loss, RNase H1-overexpressing B cells (referred to as H1 ^Tg/Tg^) were able to proliferate normally upon stimulation and did not show overt defects in class switch recombination (Maul *et al*., 2017).

We first assessed the impact of RNase H1 overexpression on the B cell transcriptome using RNA-seq and EU-seq datasets. RNase H1-overexpressing resting B cells showed significant transcriptome differences characterized by the premature expression of numerous pro-growth genes such as cell cycle and DNA repair-related genes that are usually only induced post-stimulation (Figure 5A, B). Signs of an induced innate signaling response were also observed, along with a reduced adaptive immune response. When focusing on the IgH locus, we observed premature induction of several immunoglobulin genes, in particular *Igha* (Figure 5C). This was also observed when analyzing nascent transcription using EU-seq. This surprising effect was, however, not attributable to an erroneous premature induction of the entire B cell stimulation program, as key stimulation genes such as *Nfil3* or *Aicda* were not induced in resting B cells and only became expressed post-stimulation (Figure S4A, B). It therefore appears that RNase H1-overexpressing resting B cells undergo an unusual type of premature differentiation towards the stimulated state, despite the lack of stimulation. The gene expression differences observed between control and H1^Tg/Tg^ B cells at the resting stage were still observable post stimulation (Figure S4C, D). The differences were, however, blunted, as evidenced by a reduction in the number of differentially expressed (DE) genes and the restoration of near-normal gene expression patterns at the IgH locus (Figure S4E). This confirms that RNase H1 overexpression did not impede the induction of the B cell transcriptional program, and that differences observed at the resting stage were at least partially corrected upon stimulation.

**Figure 5:**
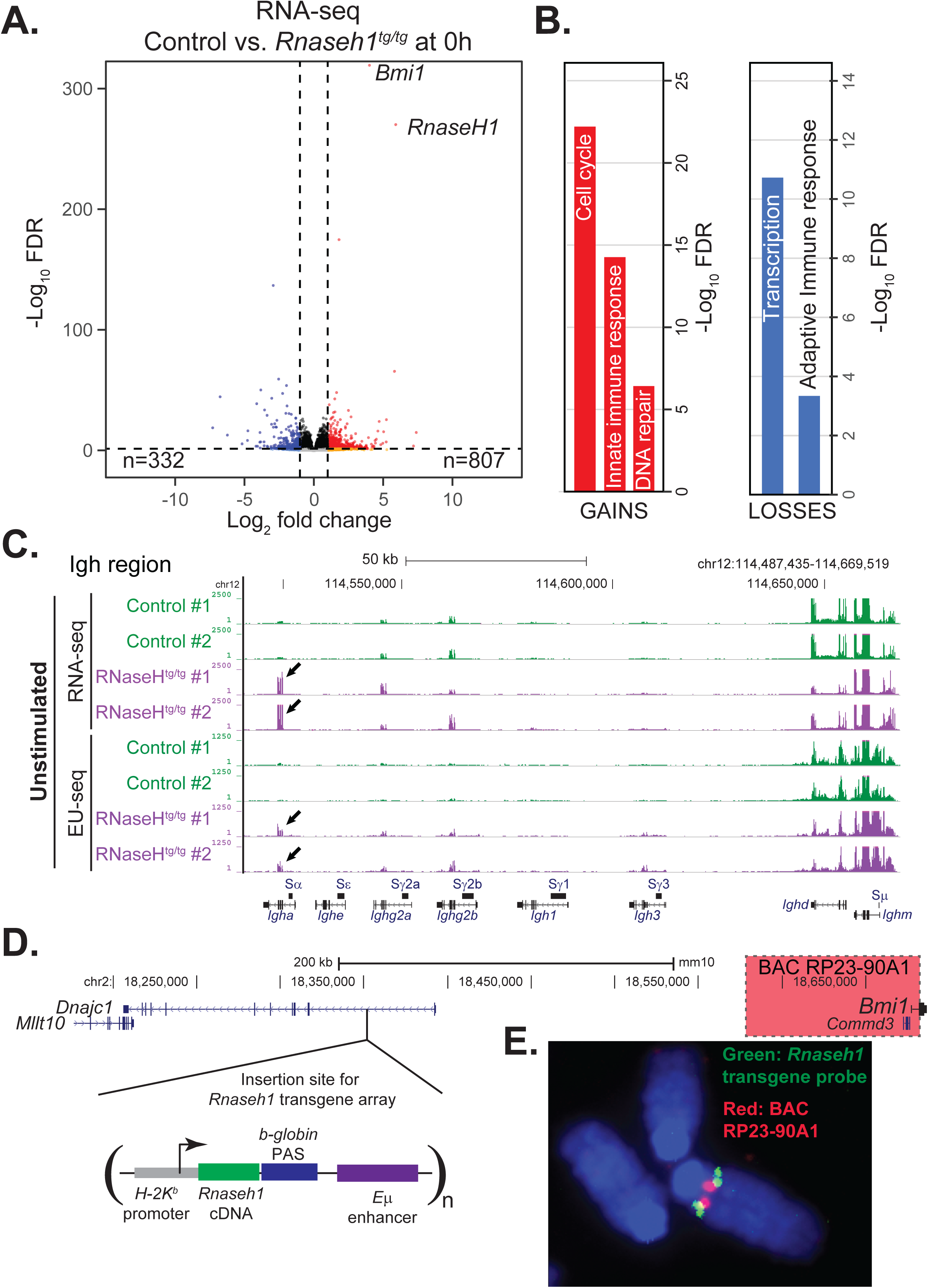
**A.** Volcano plot comparing RNA-seq data between control and *Rnaseh1*-overexpressing (*H1* ^tg/tg^) cells at the resting stage. The number of significant differentially expressed (DE) genes is indicated. The *Rnaseh1* and *Bmi1* genes are noted. **B.** Significantly enriched Gene Ontologies (David) are shown for DE genes identified in panel A. **C.** Genome browser screenshot showing RNA-seq data (top) and EU-seq data (bottom) over the IgH region showing significant overexpression of *Igha* genes (arrows) in transgenic RNase H1-expressing animals. Data is from resting (unstimulated) cells. **D.** Mapping the transgene integration site to the vicinity of the *Bmi1* gene. The genomic region surrounding the integration site is shown; the *Bmi1* gene is at the very right. The transgene was mapped to the first intron of *Dnajc1*. **E**. DNA FISH image of metaphase chromosome using a green probe for the *Rnaseh1* cDNA and a red probe for the RP23-90AI BAC, which overlaps the *Commd3* and *Bmi1* gene (highlighted red).

RNA-seq showed that the most significantly overexpressed gene, before *Rnaseh1* itself, was the *Bmi1* gene (Figure 5A). To understand this further, we mapped the integration site of the *Rnaseh1* transgene using both DNA FISH and comparative genomic hybridization and determined that it resided on chromosome 2 and had inserted in the first intron of the *Dnajc1* gene (Figure 5D, E). *Dnajc1* is itself neighboring the *Bmi1* gene. We therefore surmised that *Bmi1* overexpression was caused inadvertently by the nearby integration of tandem copies of the *Rnaseh1* transgene, which is driven by the powerful B cell-specific Eμ enhancer. The Bmi1 protein is a component of the PRC1 complex that regulates developmental gene expression by silencing genomic targets. BMI1 is known to regulate the cell cycle and cellular DNA repair capacity (Fitieh *et al*, 2021); its overexpression is often observed in human cancers and known to act in an oncogenic fashion to drive stem-like properties that favor proliferation (Bhattacharya *et al*, 2015). Due to this, we cannot ascribe the transcriptome changes observed in RNase H1-overexpressing B cells to RNase H1 itself and in fact suggest that they are due to Bmi1 overexpression.

### Nuclear RNase H1 overexpression does not significantly reduce R-loop levels or distribution

Despite the unexpected transcriptional and developmental consequences associated with the RNase H1 transgene, this animal model provided us with the opportunity to determine whether a 50-fold overexpression of nuclear RNase H1 would reduce co-transcriptional R-loop levels, as might be expected, or alter their genomic distributions. We therefore conducted R-loop profiling using DRIP-seq on both resting and stimulated B cells over-expressing, or not, nuclear RNase H1.

In resting B cells, R-loop loads were broadly increased in H1^Tg/Tg^ cells (Figure 6A) compared to controls. This surprising effect can be attributed at least in part to the gene upregulation noted above (Figure 5) and linked to the indirect consequences of transgenic RNase H1 overexpression. Gene ontology analysis for genes showing R-loop increases were consistent with those observed for over-expressed genes identified by RNA-seq (Figure S5A) and 34.6% of genes showing increased expression also showed increased R-loop loads (Figure S5B). In stimulated cells, we observed little to no difference in R-loop profiles between control and H1 ^Tg/Tg^ cells (Figure 6B), indicating that RNase H1 overexpression did not lead to a strong reduction of R-loop loads either in resting or in stimulated B cells. Inspection of specific genomic regions confirmed these trends. Strong R-loop peaks can be observed over IgM/Sμ region in resting cells, with clear R-loop induction over the Ighg1/Sγ1 region upon stimulation, and a tendency for extra R-loops to appear over the Ighg3/Sγ3 region in RNase H1-overexpressing cells (Figure S5C). R-loop patterns over a large 2 megabase region containing numerous well-expressed housekeeping genes confirmed the presence of consistent and robust R-loop peaks in RNase H1-over-expressing cells (Figure 6C).

**Figure 6.**
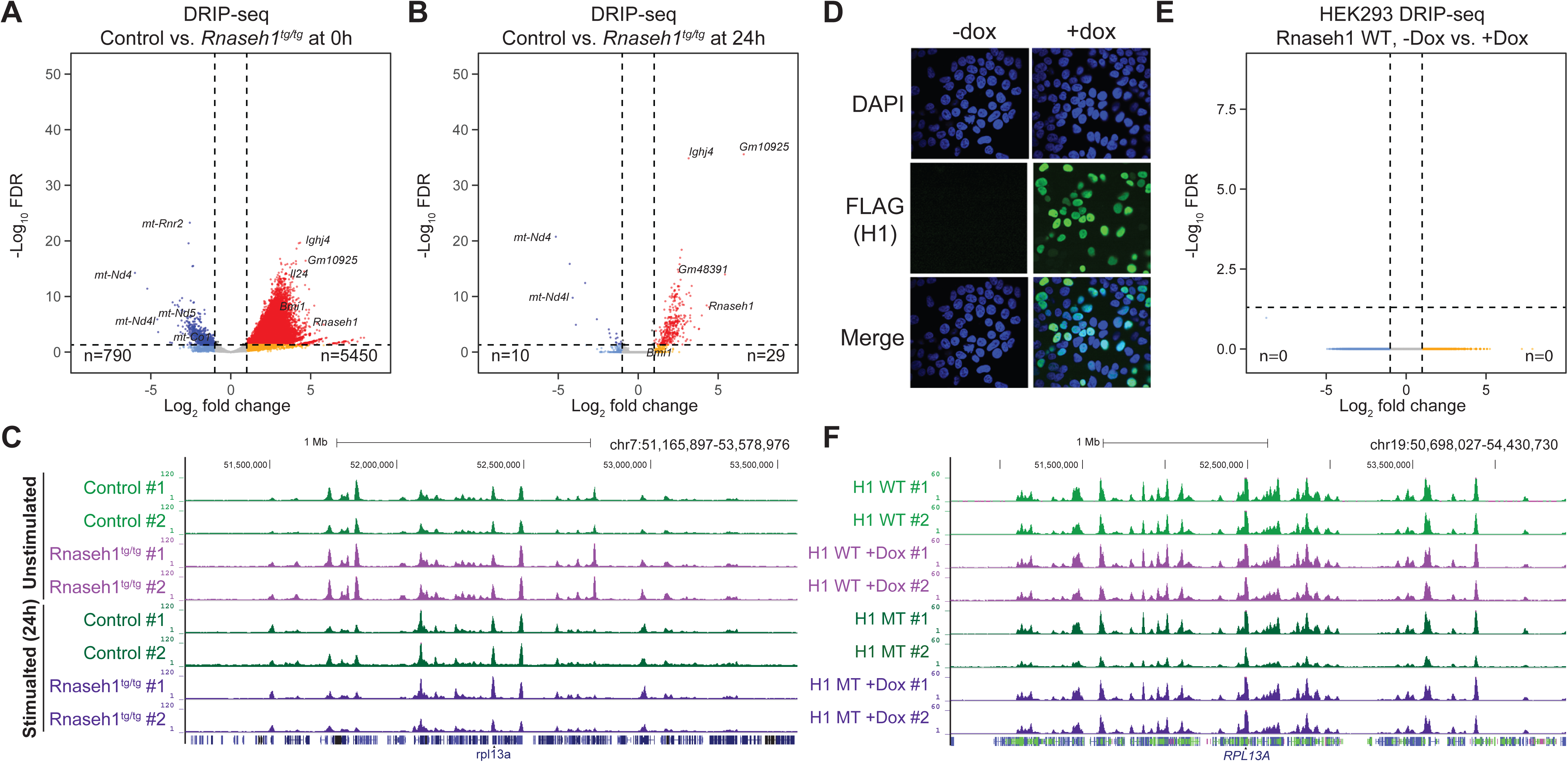
Volcano plots comparing DRIP-seq data between normal and *Rnaseh1*-overexpressing (*H1* ^tg/tg^) cells at the resting (**A**) and stimulated stages (**B**). The number of significant DE genes is indicated below. **C.** Genome browser screenshot showing R-loop distribution in resting and stimulated B cells over a ∼2 megabase region on chromosome 7 that carries many actively transcribed genes. **D.** Immunofluorescence microscopy shows inducible expression of a FLAG-tagged nuclear RNase H1 protein in HEK293T cells 24 hours after induction by doxycycline. The nuclei are marked by DAPI staining (blue). **E.** Volcano plots comparing DRIP-seq data between uninduced and induced HEK293T cells. **F.** Genome browser screenshot showing R-loop distribution in uninduced and induced HEK293T cells over a ∼2 megabase region on chromosome 19 that carries many actively transcribed genes including the *RPL13A* gene.

To independently validate these surprising findings, we designed a lentiviral, doxycycline-inducible, expression vector encoding for a FLAG-tagged nuclear RNase H1 either in an active or a catalytically inactive form (D210N) and generated stably expressing cell lines in HEK293T cells. As expected, doxycycline addition led to a time- and dose-dependent increase in RNase H1 expression (Figure S5D), and immunofluorescence microscopy clearly showed the protein was localized to the nucleus and was consistently induced in over 50% of the cell population on average, regardless of cell cycle phase (Figure 6D). Activity gel-based assays further revealed that the protein collected in cell lysates was catalytically active against labeled RNA/DNA hybrids (Figure S5E). DRIP-seq was conducted on cells treated, or not, with doxycycline and R-loop distributions were mapped. Differential gene expression analysis revealed that, consistent with the B cell data, there were no significant changes in R-loop distributions upon expression of the active RNase H1 enzyme (Figure 6E). A representative screenshot of a 2 megabase genomic region that includes numerous highly-expressed and R-loop forming genes confirmed the absence of any significant differences in R-loop distribution (Figure 6F).

## DISCUSSION

Our work reinforces the notion that RNase H1 plays a major role in mitochondrial biology, particularly in mtDNA genome replication. Early evidence showed that murine embryos lacking *Rnaseh1* failed early during development due to a loss of mtDNA (Cerritelli *et al*., 2003). Human patients carrying *RNASEH1* mutations often exhibit adult-onset muscle weakness accompanied by progressive external ophthalmoplegia due to defective mitochondrial function (Bernardino Gomes *et al*, 2024). Here, we show that loss of RNase H1 in murine B cells triggers a greater than 10-fold reduction in mtDNA copy number, accompanied by profound loss of expression of mitochondrial genes, and aberrant mitochondrial morphology. Loss of RNase H1 compromises the response to B cell stimulation and leads to failure of B cell proliferation, antibody production and the induction of the UPR^mt^ response. Interestingly, depletion of mtDNA copy number using a dominant negative mutant of the main mtDNA replication helicase, Twinkle (Milenkovic *et al*, 2013) resulted in increased expression of Atf4 and UPR^mt^ induction (Kuhl *et al*, 2017). Similarly, an astrocyte-specific knockout of Twinkle also exhibited UPR^mt^ activation (Ignatenko *et al*, 2023). These findings are consistent with our study and indicate that loss of mtDNA triggers the UPR^mt^ stress response. Collectively, this indicates that the phenotypes observed here in B cells lacking *Rnaseh1* are due to the defect in mtDNA replication.

Studying a possible non-mitochondrial role for RNase H1 based on the analysis of RNase H1 loss of function remains difficult due to the severity of mitochondrial defects. One possible non-mitochondrial function was proposed while studying a liver-specific conditional *Rnaseh1* knockout in mouse. As expected, mitochondrial function and morphology were greatly affected, but liver cells lacking RNase H1 failed to respond to antisense DNA-like oligonucleotide treatment (Lima *et al*., 2016), suggesting a nuclear role for RNase H1 in cleaving pre-mRNA:DNA hybrids. R-loops that form co-transcriptionally are another proposed substrate for RNase H1 and purified RNase H1 efficiently resolves R-loops *in vitro*. We show here that loss of RNase H1 in murine B cells had little measurable impact on the distribution or frequency of R-loop structures in the nuclear genome. This suggests that RNase H1 does not play a major role in R-loop resolution in murine B cells. It is possible that this function is instead performed by the Ribonuclease H2 complex or by a variety of other RNA/DNA helicases that have been implicated in R-loop metabolism (Brickner *et al*, 2022).

To overcome the limitations associated with loss of RNase H1 function, we used instead a B-cell-specific transgenic system that achieves a 50-fold increased expression of nuclear RNase H1 for which enzyme activity was verified in cell lysates (Maul *et al*., 2017). While this particular transgene was associated with unexpected transcriptional perturbations, it nonetheless provided an opportunity to measure R-loop distributions genome-wide. Surprisingly, R-loop levels and distributions appeared unaffected by RNase H1 overexpression. A similar observation was made from human HEK293T cells expressing an inducible nuclear RNase H1 shown to be enzymatically active in lysates.

These findings are surprising given the well-documented ability of RNase H1 overexpression to suppress genome instability cellular phenotypes (Cerritelli *et al*., 2022) such as slow replication fork speed or replication stress, the appearance of DNA damage markers such as γH2AX or 53BP1, or the appearance of DNA double-stranded breaks themselves. The genome-stabilizing effects of RNase H1 overexpression have been reported across a vast array of maladaptive conditions including hyper-transcription (Gorthi *et al*, 2018; Kotsantis *et al*, 2016; Stork *et al*, 2016), defective RNA splicing, packaging, or export (Huertas & Aguilera, 2003; Luna *et al*, 2024; Paulsen *et al*, 2009), dysfunctional topological control (Duardo *et al*, 2024; Promonet *et al*, 2020; Tuduri *et al*, 2009), altered chromatin maintenance (Bayona-Feliu *et al*, 2021; Bayona-Feliu *et al*, 2017), and others. Given the biochemical activity of RNase H1 against RNA/DNA hybrids, this has been widely interpreted to indicate that the various genome instability phenotypes were caused by the accumulation of harmful co-transcriptional R-loops.

One possibility to reconcile our findings with the dominant thinking is to suggest that RNase H1 acts predominantly or exclusively on populations of genotoxic R-loops that arise under maladaptive conditions. Under this notion, RNase H1 would not have a major role in R-loop metabolism under basal conditions. It is possible, for instance, that RNase H1 may become activated as part of the DNA damage response. It is also possible that “harmful” R-loops may possess properties that are intrinsically distinct from those of R-loops that form under basal conditions, making them more prone to RNase H1 action. While genotoxic R-loops have been often invoked, their characteristics and distributions have seldom been established at the genomic level. It will be critical for future studies to carefully define genotoxic R-loop populations using adequate genomics approaches. Supporting claims that DNA damage phenotypes arise from harmful co-transcriptional R-loops will further require demonstration that such R-loops respond to RNase H1 overexpression and that harmful R-loop loci correspond to sites of DNA breakage or fork stalling. A second, not mutually exclusive, possibility to reconcile our findings is that RNase H1 may have regulated access to distinct R-loop classes. This notion arose earlier from observations that RNase H1-based and S9.6-based R-loop maps were only partially concordant (Chen *et al*., 2017), suggesting that RNase H1 may be restricted to gene promoters, terminal regions, and enhancers (Castillo-Guzman & Chedin, 2021; Miller *et al*., 2022). Given this, the fact that RNase H1 overexpression didn’t reduce elongation-associated R-loops in our study may not be entirely surprising as these tend to not be bound by RNase H1. We note, however, that no effect on R-loop levels was noted over promoter and terminal gene regions, even though these tend to represent consensus R-loop regions (Miller *et al*., 2022). Our study therefore raises the possibility that co-transcriptional R-loops may not be sensitive to RNase H1 activity. Alternatively, it is possible that the levels of RNase H activity even under overexpression conditions are insufficient to efficiently manage the range of endogenous RNA/DNA hybrid substrates. In either case, this would suggest that the ability of RNase H1 to suppress genome instability phenotypes may be separated from its ability to resolve co-transcriptional R-loops. If confirmed, this would cause a significant re-evaluation of current models involving genotoxic R-loops.

It remains possible that RNase H1 possesses novel, as-yet-unknown or poorly described, RNA/DNA hybrid substrates that connect directly to genome instability phenotypes known to be suppressed by overexpression. Recent work has identified a type of replication-fork associated RNA/DNA hybrids (RF-RDs) (Song *et al*, 2025; Xu *et al*, 2025) that are enhanced at stalled forks and require processing to permit fork restart. Similarly, post-replicative RNA/DNA hybrids were observed behind active replication forks under conditions of transcription-replication conflicts (Stoy *et al*, 2023) and RNase H-catalyzed degradation of such hybrids was necessary for postreplicative fork repair (Heuze *et al*, 2023). Interestingly, a role for nuclear RNase H1 in supporting DNA replication forks would be consistent with its well-known role in enabling mitochondrial DNA replication by removing primers from the DNA template (Holmes *et al*., 2015). Future work will determine if such hybrids represent the long sought-after RNase H1 substrates that connect RNA metabolic processes to DNA-based genome instability phenotypes.

## METHODS

### Mouse strains

All mouse strains were bred and housed in accordance with National Institutes of Health and United States Public Health Service policies. Animal research was performed only after protocols were approved by the *Eunice Kennedy Shriver* National Institute of Child Health and Human Development Animal Care and Use Committee. Previously, we described cell lines derived from a hydroxytamoxifen-inducible Cre mice carrying floxed *RNaseh1* alleles engineered to contain loxP sites flanking exons 5 and 7 (Holmes *et al*., 2015). These mice were crossed with the *mb1-Cre* mouse to produce a conditional *Rnaseh1* deletion mouse model in which the *Rnaseh1* gene is knocked out at the early pro B cell stage. *Mb1-Cre* mice were obtained from Dr. Herbert Morse and used with the permission of Dr. Michael Reth (Hobeika *et al*., 2006). Genotyping was conducted by PCR on DNA isolated from mouse tail clips using the Qiagen DNeasy Blood and Tissue kit according to the kit’s instructions. PCR products were analyzed on agarose gels to confirm genotypes. Direct lysis protocol from Viagen was also used to obtain mouse tail DNA. Mouse genotyping has been more recently outsourced to Transnetyx Inc. PCR primers used for genotyping are listed in **Table S2**. The RNase H1 overexpression model (H1^tg/tg^) in B cells was previously described (Maul *et al*., 2017).

### B cell collection, *ex vivo* stimulation, and Western blots

Following euthanasia, we extracted the spleen from H1^fl/fl^, H1^fl/del^, H1^del/del^, H1^tg/tg^ and control H1^wt/wt^ mice. Single-cell suspensions of splenic cells were prepared in PBSS (PBS+3% FBS). Splenic cell pellets were treated with ACK lysis buffer first, spun down and then resuspended in MACS buffer (PBS pH 7.2, 0.5% BSA, 2 mM EDTA). After filtering through 40 µm cell strainers, cells were spun again, and resting B cells were isolated from these pellets by depletion of non-B cells using magnetically labeled biotin-conjugated antibodies and anti-biotin microbeads from mouse B cell isolation kit (Miltenyi Biotec). The total number of splenic cells present in each cell pellet and purified B cells were counted using a cell counter (Cellometer Auto T4 – Nexcelcom Biosciences). The purity of isolated B cells was checked by staining the cells with fluorochrome-conjugated antibody CD45R (B-220)-FITC and Anti-Biotin-APC and analyzed with Flow cytometry using a BD Accuri C6 Flow Cytometer. Purified B cells were cultured in suspension at 37°C and 5% CO_2_ using RPMI media with 10% (v/v) FBS, 2 mM L-Glutamine, 100 U/ml Penicillin-Streptomycin, and 100 μM beta-mercaptoethanol. Cells were plated at 1 x 10^6^ cells/ml and *ex vivo* stimulation was done with 20 μg/ml LPS (Lipopolysaccharide; *E. coli* serotype 0111: B4) and 25 ng/ml recombinant cytokine IL4 (Interleukin-4) for 1-5 days. Aliquots were taken from cultures at 24, 48, 72 and 96 hours and cells were counted. Number of cells present at every time points were plotted for H1^fl/fl^, H1^fl/del^ and H1^del/del^ B cell cultures to check for continuity of proliferation. RNase H1 hybrid enzymatic activity was measured using the gel renaturation assay using 15% SDS polyacrylamide gel which contained uniformly labelled poly-^32^P-rA/poly-dT substrate (Crouch *et al*, 2001). We varied the number of B cells between 0.1 to 0.5 million cells for H1^fl/del^ and 0.5 to 2 million cells for H1^del/del^. To check for circulating IgG in the blood of H1^fl/del^ and H1^del/del^ mice, the serum from these mice were mixed with equal amount of 2xSDS-PAGE buffer and a western was performed using anti-mouse IgG antibody (Pierce) and ECL prime western blotting reagent (GE healthcare).

### Mitochondrial copy number determination

DNA from resting and 24 hours stimulated H1^fl/fl^, H1^fl/del^ and H1^del/del^ B cells were extracted using Puragene DNA kit according to kit’s instruction. Quantitative PCR analysis was performed using a Lightcycler 480 and SYBR Green 1 Master reagent. Gene copy numbers were calculated as percentage of wild type. Primers used to check genomic *Rnaseh1* gene and mitochondrial DNA copy numbers are listed in **Table S2**. For RT-qPCR, total RNA was extracted from resting and stimulated B cells using Trizol reagent, PreLink DNase and PureLink RNA Mini Kit. RNA was also prepared using Maxwell 16 LEV simplyRNA Cells Kit for the validation of RNA-seq results of few genes. Extracted RNA was reverse transcribed with Omniscript RT kit and oligo-dT primers into cDNA. Expression levels of target genes were normalized to the *ACTB* housekeeping gene. Primers for all the genes analyzed via RT-qPCR are listed in **Table S2**. Westerns were performed using about three million B cells which were lyzed in RIPA buffer with protease / phosphatase-inhibitor mini tablets (Invitrogen) for 30 min on ice. SDS-PAGE gels (10-20%) were loaded with equal number of B cells and after electrophoresis, proteins were transferred to PVDF membranes via wet transfer in Towbin’s transfer buffer. Membranes were blocked with Azure chemiluminescent blocking buffer at RT for 1 h and then incubated with diluted antibodies (1:1,000 for Primary antibody and 1:10,000 or 1:20,000 for fluoro/chemi secondary antibodies). Azure fluorescent washing buffer was used for washings and ECL substrate was used to develop western blots.

### Electron microscopy

Resting and stimulated H1^fl/del^ and H1^del/del^ B cells were collected and fixed at room temperature with 2.5% glutaraldehyde in 0.1 M sodium cacodylate buffer and centrifuged into a pellet. Cell pellets were then put into 2% Agarose made in 0.1 M sodium cacodylate buffer, pH 7.4. The following processing steps were carried out using the variable wattage Pelco BioWave Pro microwave oven (Ted Pella, Inc., Redding, CA). Cells were rinsed in 0.1 M sodium cacodylate buffer, pH 7.4, post-fixed in 1% osmium tetroxide made in 0.1 M sodium cacodylate buffer, rinsed in double distilled water (DDW), 3% (aq.) uranyl acetate enhancement, DDW rinse, ethanol dehydration series up to 100% ethanol, followed by a Embed-812 resin infiltration series up to 100% resin (Electron Microscopy Sciences, Hatfield, PA). The epoxy resin was polymerized for 20 hours in an oven set at 60° C. Ultra-thin sections were prepared on a Reichert-Jung Ultracut-E ultramicrotome (80 nm). The cooper grids were post-stained with uranyl acetate and lead citrate and examined in a JEOL 1400 transmission electron microscope operating at 80kV and images were acquired on an AMT BioSprint 29 camera. ImageJ was used to measure and analyze these images for total cell, nuclear, and cytoplasm areas.

### Transcriptome analysis by RNA-seq and EU-seq

RNA-seq analysis was performed for H1fl/del, H1del/del, H1tg/tg and control H1wt/wt B cells at the resting stage and 24 hours post stimulation. RNA was collected using Maxwell 16 LEV simplyRNA Cells Kit and RNA purity was checked by obtaining RNA integrity number via Bioanalyzer experiment using Agilent RNA 6000 Nano Kit. RNA-seq library construction and sequencing was performed using NIH Intramural Sequencing Center (NISC) services for biological triplicates of RNAs extracted from resting and 24 hours-stimulated H1^fl/del^ and H1^del/del^ B cells. Molecular Genomics Laboratory Core Facility of NICHD performed RNA-seq library construction and sequencing for eight samples each of resting and 24 hours stimulated H1^tg/tg^ B cell RNA, and six samples each of resting and 24 hours stimulated control H1^wt/wt^ B cell RNA. Nascent transcripts were labeled with 5-ethynyl uridine (EU) (0.5 mM final concentration) for 20 min prior to harvesting. Total RNA was isolated and EU-RNA was enriched using Click-iT Nascent RNA Capture Kit (Invitrogen), with 5 µg of EU-RNA for biotinylation by click reaction and 1 µg of biotinylated RNA for binding to Streptavidin T1 magnetic beads. cDNA was generated using iScript Select cDNA Synthesis Kit (Bio-Rad). cDNA was cleaned up before second-strand synthesis. Then cDNA was ligated to Illumina barcoded adapters for sequencing. Library quality was checked on an Agilent BioAnalyzer and sequencing were performed on Illumina HiSeq or NovaSeq instruments.

### R-loop mapping by DRIP-seq

B cells at the resting and stimulated (24 h) stages were purified for H1^fl/del^, H1^del/del^, H1^tg/tg^ and control H1^wt/wt^ strains. Cells were lysed overnight in lysis buffer and processed for S9.6-based R-loop mapping using the DRIP-seq protocol or, alternatively, the high-resolution, strand-specific sDRIP-seq protocol (Sanz *et al*., 2021).

### Computational analysis

Fastq files were aligned to the mouse (mm10) or human (hs1/T2T) genomes using STAR-aligner v2.3 (https://github.com/alexdobin/STAR) and filtered for concordant (paired-end) and uniquely mapped reads using samtools to generate BAM files. To generate bigWig files for EU-seq, DRIP-seq and sDRIP-seq, BAM files were further duplicate-removed using samtools rmdup. BAM files were processed into bedGraph files using bedtools genomecov, normalized against total mapped and filtered reads. BedGraph files were turned into bigWig files using wigToBigWig. To generate regions used in DESeq2 statistical analysis for EU-seq and DRIP-seq/sDRIP-seq, peaks were called using macs2 (2.2.9.1) using “ --nolambda --nomodel”. Peaks were merged from each sample group (EU-seq, DRIP-seq, and sDRIP-seq). For DRIP-seq, we used restriction digest regions that overlap with the macs2-called peaks above (RE digest: HindIII, EcoRI, BsrGI, XbaI, and SspI), filtering out low count peaks (baseMean < 20). For RNA-seq, we used gencode mm10 annotation with the highest APPRIS principal. htseq-count (2.0.0) was used to count mapped reads from BAM files (not duplicate removed) onto the regions described above. Finally, DESeq2 was performed to find significantly increased/decreased regions and genes (padj <= 0.05, log2FC +/-1). DAVID Gene Ontology (DAVID Dec 2021, DAVID Knowledgebase v2023q4) was used to find significant GO Terms of these genes.

### Localization of transgene insertion site

Metaphase chromosomes were prepared from H1^tg/tg^ transgenic mouse tumor cell lines by adding 0.075 M KCl to Colcemid-treated cells. Metaphase chromosomes were then karyotyped using DAPI Banding and FISH analysis was performed with an *Rnaseh1* transgene DNA probe and wild-type chromosome 2 BAC plasmid (RP 23-90 A1 with *Bmi1* gene) DNA probe. Comparative genomic hybridization studies were performed using H1^tg/tg^ transgenic cell line DNA and wild-type kidney DNA. DNA were prepared using Puregene Core Kit and digested with AluI and RsaI, purified, labeled with Cyanine 3 (Cy3-green) for cell line DNA and with Cyanine 5 (Cy5-red) for wild-type Kidney DNA. Cell line and wild-type DNA were mixed in equimolar ratio, hybridized to a slide with probes printed and read through Confocal scanner. The ratio calculated by the software showed that there was a gene deletion (*Dnajc1*) in chromosome 2 of the transgenic cell line. The exact 3’-end of transgene insertion site was determined by sequencing PCR fragment using forward primer from *Rnaseh1* transgene (5’-GCC TCA CCT CAG AGA AAC CAG ACA-3’) and reverse primer from wild-type chromosome 2 at position 18,353,100 bp (5’-AAT GTG GGA GGA AGC CAG GAG G-3’).

### Inducible nuclear RNase H1 expression in HEK293T cells

We cloned the cDNA for the nuclear form of human RNASEH1 (excluding the first 27 amino acids) into pLVX-TetOne-Puro 3xFLAG (a kind gift from Dr. Priya Shah’s laboratory, UC Davis), a lentiviral vector that permits Dox-inducible expression. The cDNA was PCR amplified using primers introducing unique ClaI and BamHI sites and recloned after digestion of pLVX-TetOne-Puro with ClaI and BamHI (New England Bio Labs, MA, USA). Site-directed mutagenesis was used to introduce the D210N mutation in the enzyme’s active site to create a catalytically inactive version of the protein. The pLVX-TetOne-Puro vector carrying human RNase H1 WT or mutant genes were named pLVX-TetOne-Puro-hRNASEH1 WT-3xFLAG and pLVX-TetOne-Puro-hRNASEH1 Mut-3xFLAG, respectively. For lentivirus production, the lentiviral plasmids (5 μg), packaging, and envelope plasmids (4 μg of psPAX2 and 1.5 μg of pMD2.G) were co-transfected into 70% confluent HEK293T cells in 10 cm culture dishes using TurboFect Transfection Reagent (Thermo Scientific). The medium was replaced 16 h later. The viral supernatant was collected 24 h after changing medium, filtered through a syringe with 0.45 μm filters to remove cell debris and 1 mL of it was used to infect HEK293T cells in a 6-well plate in the presence of 7 µg/mL polybrene (Sigma-Aldrich). The medium was replaced 20 h later. After 48 h of infection, cells were selected with 0.55 μg/mL of Puromycin (Sigma-Aldrich) for 7 days. The puromycin-resistant cells were then expanded. HEK293T cells stably transduced with lentiviral vectors were induced with 100 μg/ml doxycycline (+Dox) or vehicle (+DMSO) for 24-48h in DMEM with 10% Tet-system approved FBS (Takara Bioscience).

HEK293T cells were cultured at 37°C in 5% CO2 in DMEM supplemented with 10% FBS and 1% penicillin/streptomycin. Cells were regularly tested and confirmed to be negative for mycoplasma before experiments. Samples were seeded with cells at equal densities 1–2 d before experiments and were grown to ∼50–70% confluence. Human RNASEH1 overexpression was induced by the addition of doxycyline to growth media. Media and puromycin were replaced every 1-3 days. RNase H1 overexpression was confirmed by qRT-PCR, Western blotting, or immunofluorescence microscopy using the anti-FLAG M2 primary antibody (F1804; Sigma-Aldrich). Enzymatic activity was measured using gel renaturation assays, as described (Crouch *et al*., 2001).

## Acknowledgements

This work was funded by the National Institutes of Health grant R35 GM139549 (to FC) and by the Division of Intramural Research at the *Eunice Kennedy Shriver* National Institute of Child Health and Human Development National Institutes of Health (to RJC and SMC).

## Data Availability

All NextGen sequencing files were deposited on the NCBI GEO repository under accession number GSE292947 (accessible with private reviewer token ybejiygcbxyxjct).

## Supplementary Materials

**Figure S1:**
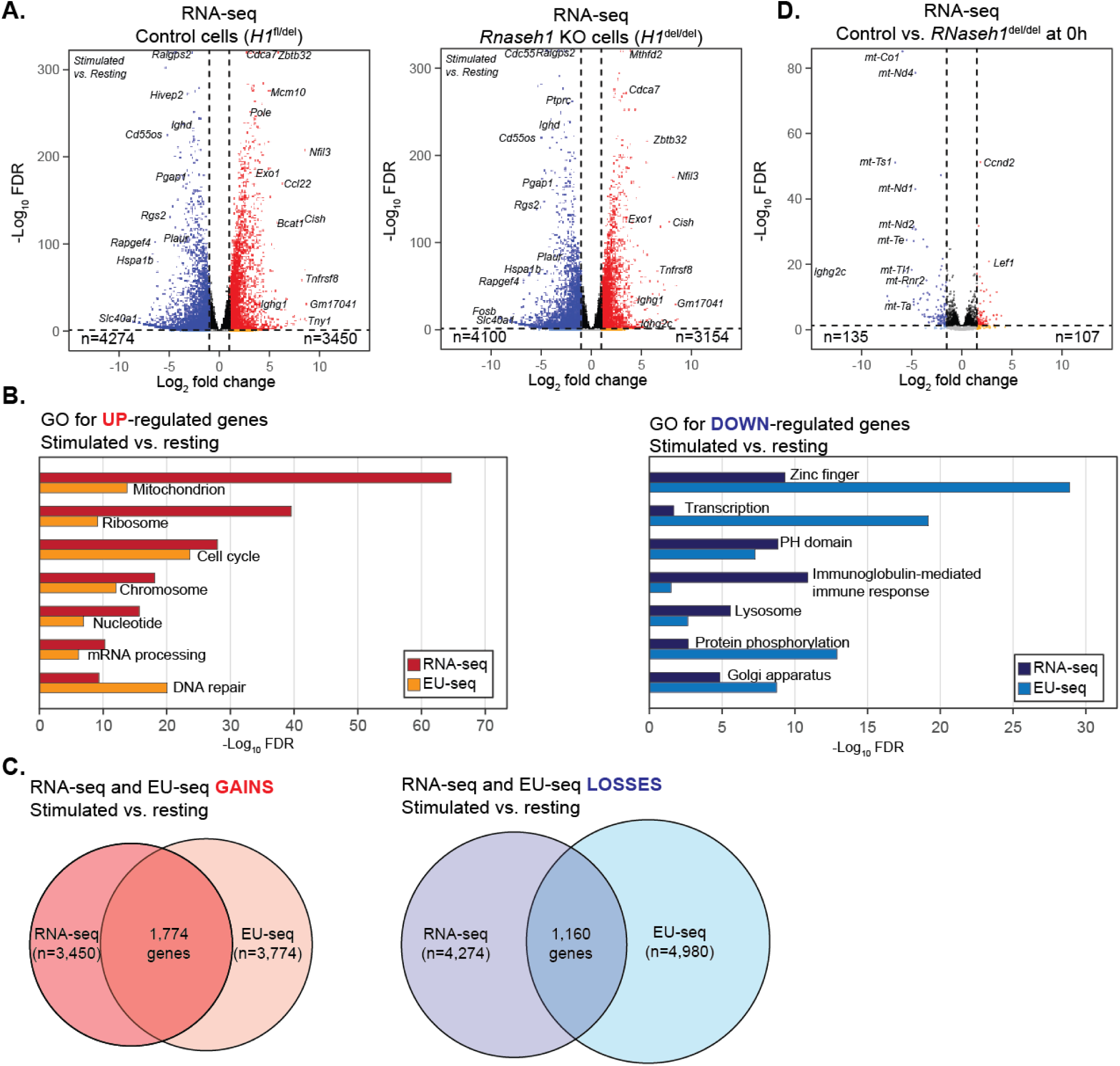
**A.** Volcano plots of RNA-seq data comparing gene expression differences between resting and stimulated B cells in both control cells (*H1* ^fl/del^) and *Rnaseh1* KO cells (*H1* ^del/del^). Vertical dashed lines indicate |Log_2_FC|>1 and the horizontal dashed line indicates a Log_10_(FDR)<0.05. The total numbers of significantly up- or down-regulated genes are indicated. **B.** Significantly enriched Gene Ontologies (David) for up- and down-regulated genes (RNA-seq and EU-seq) comparing resting and stimulated B cells in control cells (*H1* ^fl/del^). **C**. Venn diagrams depicting overlap between RNA-seq and EU-seq data for genes that were upregulated (Gain) or downregulated (Loss) upon B cell stimulation in control cells (*H1* ^fl/del^). **D**. Volcano plot of RNA-seq data comparing gene expression differences between control cells (*H1* ^fl/del^) and *Rnaseh1* KO cells (*H1* ^del/del^) at the resting stage.

**Figure S2:**
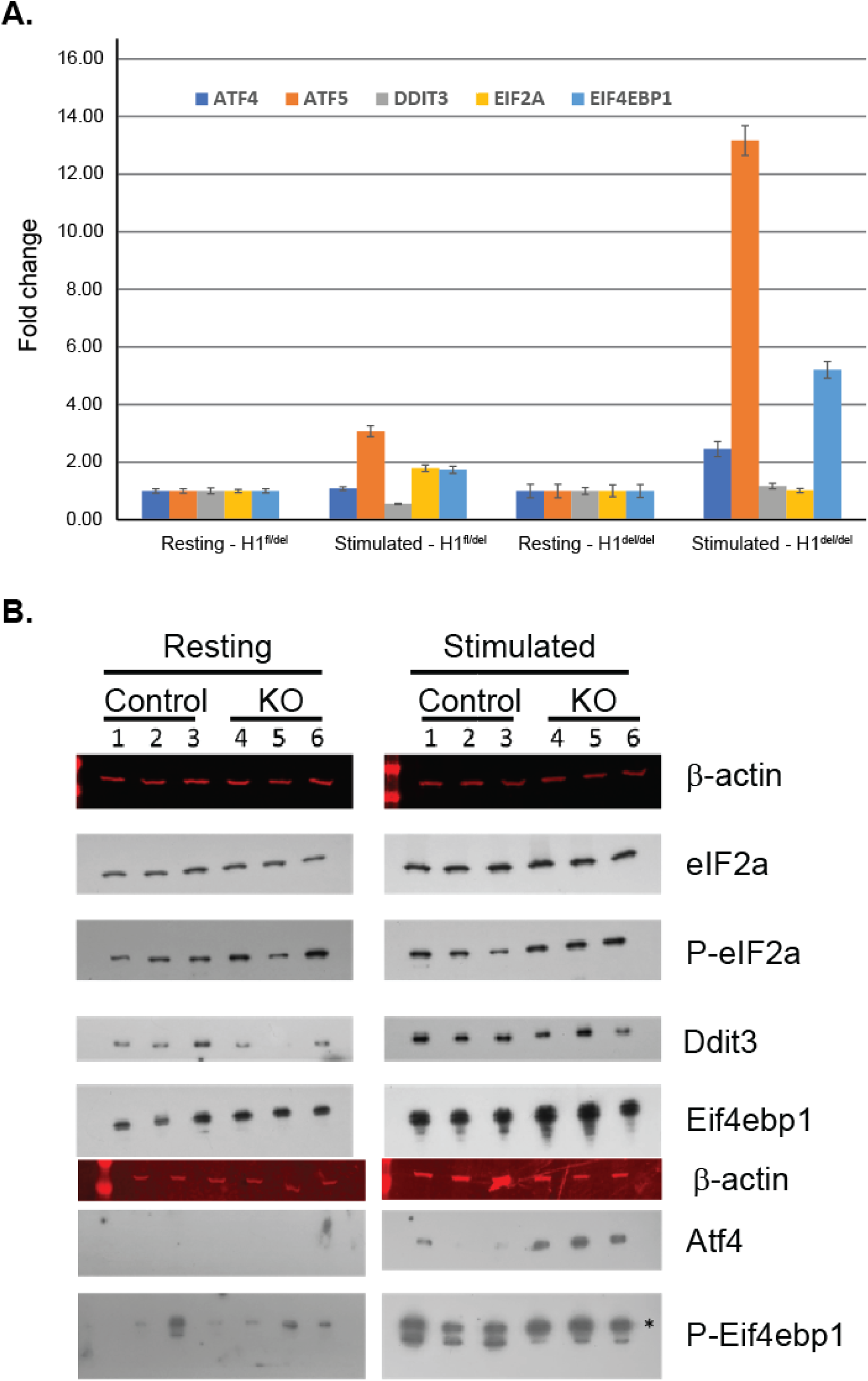
**A.** RT-qPCR validation of the UPR^mt^ induction for key genes. Data is normalized the resting B cells. Significant induction is observed for Atf4, Atf5, and Eif4ebp1. Data is from RNA extracted from three independent animals. **B.** Western blot validation for the UPR^mt^ induction. Each group displays signal from B cells extract from 3 independent animals either resting or 24 hours post-stimulation. Control group refers to *H1* ^fl/del^ mice, while KO group refers to *H1* ^del/del^ animals. The detected proteins in each panel are indicated on the right. * indicates the phosphorylated form of Eif4Ebp1. Significant increases are seen for Atf4 and P-Eif4Ebp1, with a minor increase for P-Eif2a.

**Figure S3:**
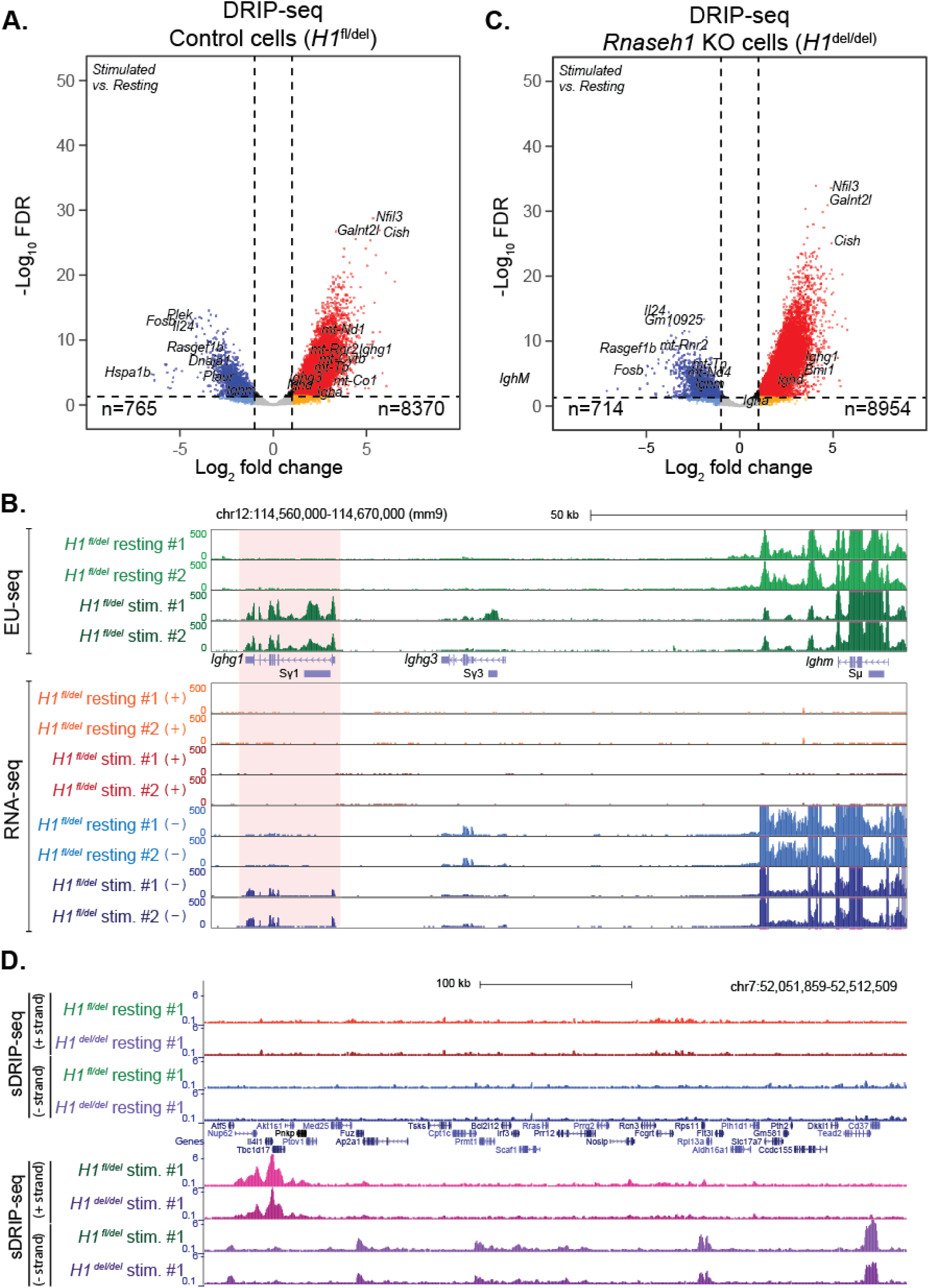
**A.** Volcano plots of DRIP-seq data comparing R-loop intensities between resting and stimulated cells for control (*H1* ^fl/del^) cells. Vertical dashed lines indicate |Log_2_FC|>1 and the horizontal dashed line indicates a Log_10_(FDR)<0.05. The number of loci that are significantly up- or down-regulated are indicated. **B.** Genome browser screenshot showing nascent transcription (EU-seq, top) and RNA-seq (bottom) over the IgH region for control samples at the resting and stimulated stages, as indicated. The IgG1 region is highlighted. **C.** As with panel A but for resting and stimulated *H1^del/del^* B cells. **D.** Genome browser screenshot over a representative autosomal region depicting strand-specific sDRIP-seq data in both control and KO cells at the resting (top) and stimulated (bottom) stages. Both cell types show similar R-loop induction patterns.

**Figure S4.**
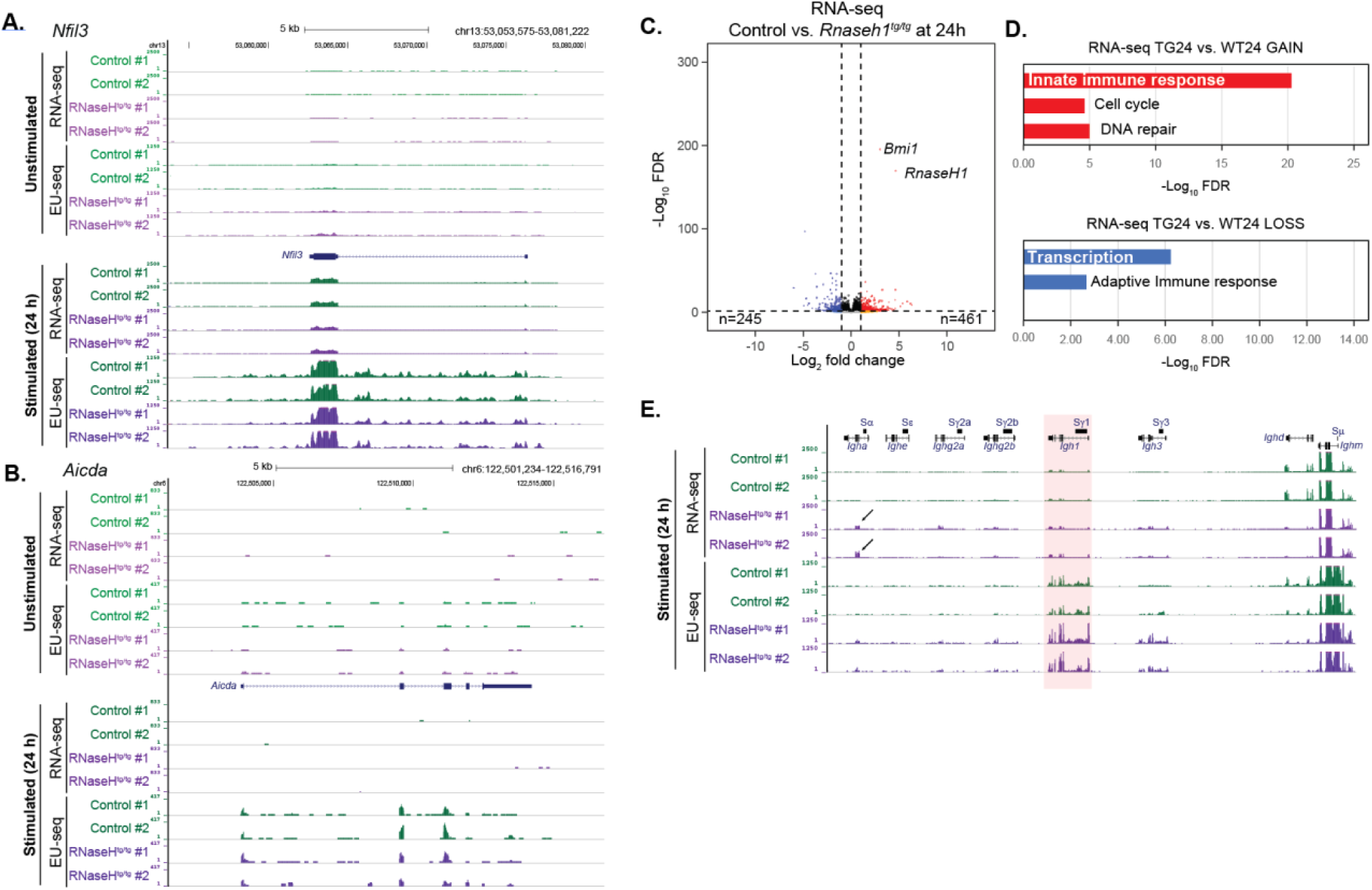
Genome browser screenshots showing gene expression data (RNA-seq and EU-seq) for the *Nfil3* (**A**) and *Aicda* genes (**B**) in both control and RNase H1-overexpressing cells. **C.** Volcano plots of RNA-seq data comparing gene expression levels between control and RNase H1-overexpressing cells 24 hours post-stimulation. **D.** Enriched GOs observed when comparing gene expression patterns (RNA-seq) between control and RNase H1-overexpressing cells 24 hours post-stimulation. **E.** Genome browser screenshot showing gene expression data (RNA-seq and EU-seq) over the IgH region for control and RNase H1-overexpressing cells 24 hours post-stimulation.

**Figure S5.**
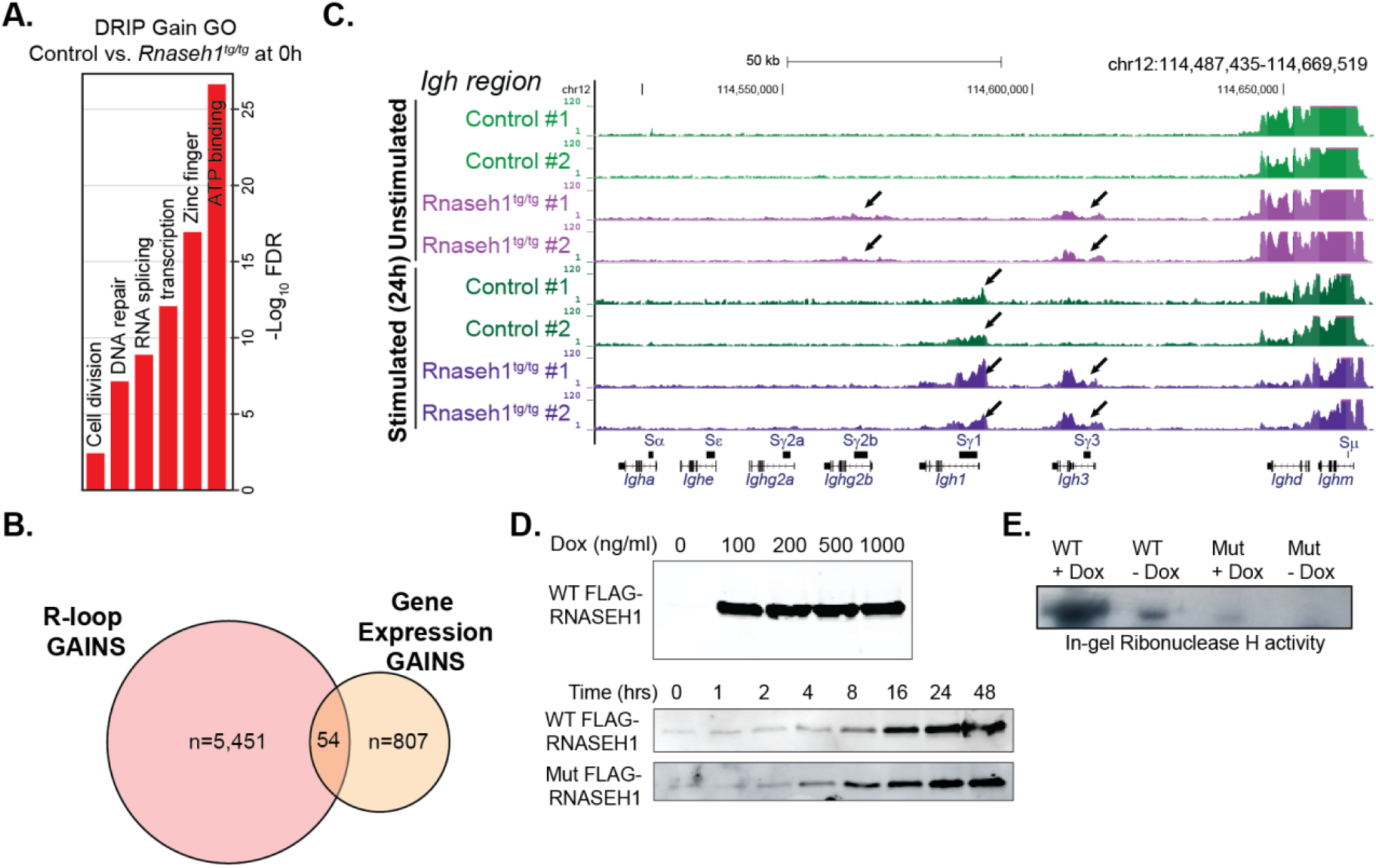
**A.** Enriched GOs observed when comparing R-loop data (DRIP-seq) between resting control and RNase H1-overexpressing cells. **B.** Venn diagrams depicting overlap between genes showing R-loop gains and gene expression gains when comparing resting control and RNase H1-overexpressing cells. **C.** Genome browser screenshot showing R-loop distribution over the IgH region for control and RNase H1-overexpressing cells at both the resting and stimulated stages. **D.** (top) Western blot showing induction of inducible nuclear RNase H1 expression in HEK293T cells as a function of doxycycline (Dox) addition; samples were collected 24 hours post induction. (bottom) Western blot showing time-dependent induction of RNase H1 expression in HEK293T cells post Dox induction (100 ng/ml). **E.** Representative RNase H1 gel renaturation assay showing RNase H1 activity against labeled RNA/DNA hybrids in lysates generated from an equal number of HEK293T cells expressing, or not, the wild-type or mutant RNase H1 protein.

